# “Positive and negative plant-soil feedbacks are caused by mycorrhizal type, soil fertility, and phylogenetic relatedness in a mixed Dipterocarp rain forest”

**DOI:** 10.1101/399295

**Authors:** R. Max Segnitz, Sabrina E. Russo, Stuart J. Davies, Kabir G. Peay

**Affiliations:** Department of Biology, Stanford University, Stanford, CA 94305-5020, USA; School of Biological Sciences, University of Nebraska, Lincoln, NE 68588-0118.; Center for Tropical Forest Science, Smithsonian Institution Washington, D.C. 20013-7012, USA

**Keywords:** tropical forest, seedling, plant-soil feedback, soil fertility, resource availability

## Abstract

While work in temperate forests suggests that there may be consistent differences in plant-soil feedback (PSF) between plants with arbuscular and ectomycorrhizal associations, it is unclear whether this is compatible with the high diversity of tropical rainforests. To examine this, we tested the effect of mycorrhizal type, phylogenetic distance, and soil fertility on variation in PSF strength in a mixed-tropical rainforest with a uniquely high diversity of ectomycorrhizal and arbuscular mycorrhizal trees. We found positive phylogenetic PSFs for ectomycorrhizal tree species that were insensitive to soil fertility. By contrast, PSFs for arbuscular mycorrhizal tree species were negative, and increasingly so with greater soil fertility. Our results demonstrate consistent effects of mycorrhizal types on plant population dynamics across biomes, and help explain biogeographic variation across tropical forests, such as familial dominance of the Dipterocarpaceae in SE Asia. However, they also raise questions about the role of PSFs in maintaining tropical diversity.

**Statement of authorship:** RMS, SER, SJD and KGP designed the experiment. RMS conducted the experiment and collected data. RMS analyzed data with input from KGP and SER. RMS wrote the first draft of the manuscript, and all authors contributed to subsequent revision and preparation of the manuscript.

## Introduction

Localized accumulation of species-specific natural enemies has been hypothesized to influence the diversity and structure of plant communities by limiting local recruitment and thereby preventing competitive exclusion by dominant species (Janzen 1970; Connell 1971). While the original Janzen-Connell framework focused primarily on herbivorous arthropods and seed predators, soil-borne microorganisms are also important natural enemies that strongly influence seedling recruitment (Augspurger 1983; Bever et al. 1997; Mangan et al. 2010a). Experiments comparing plant growth in conspecific vs. heterospecific soils often show more negative effects of microbes in conspecific soils, known as negative plant soil feedbacks (PSFs) (Bever et al. 1997; Mangan et al. 2010a; Augspurger 1983). Because of their potential role in causing conspecific negative density-dependence (CNDD), PSFs can shape plant community structure (Condit et al. 1992; van der Heijden et al. 1998; Klironomos 2002; Comita et al. 2010; Bagchi et al. 2010a; Mangan et al. 2010b; Johnson et al. 2012; Bagchi et al. 2014).

Despite the potential importance of negative PSFs in maintaining diversity in tropical forests, it is clear that the strength and direction of PSFs can vary dramatically among species and locations. For example, the same species may vary in PSF strength across sites (Liu *et al.* 2015) or range from positive to negative between co-occurring species (Bennett et al. 2016). One likely reason for this variation is that the interactions of soil-borne microorganisms with plants are complex (Peay et al. 2016). Soil microbial communities are highly diverse and include microorganisms that range from pathogens to mutualists of plants (Bronstein 1994; Johnson *et al.* 2010). In temperate forests there may be consistent differences in the benefits conferred to hosts by arbuscular mycorrhizal (AM) versus ectomycorrhizal (EM) symbioses, and thus the PSF they cause (Bennett et al. 2017). These differences may stem from variation between AM and EM fungi (AMF and EMF) in host specificity, dispersal ability, enzymatic capacities, and the interactions between soil pathogens and nutrients (Bruns & Shefferson 2004; Morris et al. 2007; Hoeksema et al. 2010). Evolving from a saprotrophic ancestor, EMF have greater enzymatic capacity to access nutrients bound in the complex organic forms found in the plant litter of nutrient-depleted soil (Read & Perez-Moreno 2003). Thus, while AM and EM symbioses are both expected to be more beneficial to plants in less fertile soil types (Hoeksema *et al.* 2010), this benefit should be greater for EM-hosting plant species. If so, then PSFs produced by AM versus EM fungi should also vary along soil fertility gradients.

Most work on tropical PSF has taken place in neotropical forests where AMF dominate as soil mutualists (Janos 1980; Torti et al. 1997), and EMF are rare or associated with monospecific stands that are considered atypical (Newbery et al. 1988; Moyersoen et al. 2001; McGuire 2007). However, EM symbiosis can be common in some tropical rainforests, for example those in Southeast Asia (Proctor *et al.* 1983; Alexander & Hogberg 1986; Brearley 2012). Because of the potentially different nature of the feedbacks produced by EM versus AM symbiosis (Fukami et al. 2017), a generalized picture of the role of PSFs in determining variation in tree species composition and maintaining diversity in tropical forests should include regions dominated by EM symbioses.

Overstory trees can shape the soil microbial community encountered by recruiting seedlings via root exudates, litter chemistry, and accumulation of host-associated mutualists and pathogens. The phylogenetic relationship between a recruiting seedling and the nearest overstory tree may influence whether these localized soil biota enhance or reduce seedling growth and/or survival (Gilbert & Webb 2007; Liu *et al.* 2011). Experiments growing seedlings in soils with different inocula have shown that PSFs are more strongly negative when seedlings and the adult tree nearest the soil inoculum source are phylogenetically closely related (Bagchi et al. 2010a; Liu et al. 2011). The causal mechanism is presumably that more closely related tree species tend to share more pathogens (Gilbert et al. 2015), although this may not be true of all plant lineages (Mehrabi & Tuck 2014). However, PSFs measured in such experiments are the net outcome of not only the negative effects caused by soil pathogens, but also the positive effects of soil mutualists, which also may be affected by evolutionary relatedness (Nara 2006; Ishida *et al.* 2007). Thus, due to the greater host specificity associated with EM symbiosis (Bruns *et al.* 2002; Davison *et al.* 2015), PSFs may also vary more strongly with evolutionary relatedness for EM, compared to AM host species.

To dissect these key sources of variation in PSF, we conducted a shadehouse experiment with seedlings of five EM and three AM tree species using a design that allowed us to quantify whether the direction and strength of PSF depends on a tree’s mycorrhizal association type, phylogenetically driven variation in the soil microbiota, and resource availability of the soil. Our study site is a high-diversity mixed dipterocarp Bornean rain forest that is co-dominated by EM and AM tree species, and in which tree species composition varies strongly along a soil fertility gradient (Davies et al 2005), allowing us to decouple the effects of these factors on PSF. We predicted that (1) PSFs in this forest would, on average, be negative, supporting negative PSF as a mechanism promoting floristic diversity, but that (2) this variation would be structured by mycorrhizal type, phylogenetic relatedness, and soil fertility. Specifically, we expected EM hosts to experience weaker negative feedbacks than AM hosts and that the strength of the interactive effects of soil type and evolutionary relatedness would be greater for EM than AM host species.

## Methods

### Study Site

Lambir Hills National Park (LHNP) is a 7,800 ha protected area in northwest Borneo in Sarawak, Malaysia (4° 20’ N, 113° 50’ E). It is classified as tropical mixed dipterocarp forest (Lee et al. 2002). A 52-ha long-term forest dynamics plot located at this site (Lee et al. 2002) is characterized by two geological formations that produce a gradient of soil texture spanning coarse, infertile sandy loam to fine, less infertile clay soil (Baillie et al. 2006). Variation in tree species composition and dynamics in the plot is strongly structured by this edaphic gradient (Davies et al. 2005; Russo et al. 2005). Over 1,200 species occur in the plot, with dominance by species of the EM Dipterocarpaceae, which account for approximately 16% of stems and 42% of basal area (Lee et al. 2002). Here, we used the well-described forest community variation along the edaphic gradient in the plot as the basis for our experimental design separating the effects of mycorrhizal type, soil type, and phylogenetic relatedness on variation in PSF.

### Experimental Design

We established a shadehouse plant-soil feedback experiment using eight focal tree species (Figure 1; Table S1; see Appendix S1). We collected seeds locally from EM and AM host trees following a general flowering event in the fall of 2013. EM hosts comprised four dipterocarp species (*Dryobalanops aromatica, Shorea curtisii, Shorea laxa*, and *Shorea scaberrima*) and one species of Fagaceae (*Castanopsis hypophoenica*). The three AM hosts (*Dacryodes expansa, Whiteodendron moultonianum*, and *Madhuca utilis*) were from three different families, Burseraceae, Myrtaceae, Sapotaceae, respectively. We grew seedlings of each species in soils collected from beneath trees of species that differed in their phylogenetic relatedness to the focal species, from conspecific, congeneric, confamilial, conordinal, or distantly related (hetero-ordinal), such that each seedling was grown in soils encompassing a range of seedling-soil phylogenetic distances. To better separate the effect of phylogenetic distance from mycorrhizal association, for each focal species, we also used both a distantly related EM host soil source and a distantly related AM host soil source (hetero-ordinal in both cases). To test the effects of soil type on PSF, for each level of phylogenetic distance, we grew seedlings in soil sourced from both sandy loam and clay. Because there are few generalist tree species that occur on both soils (Davies et al. 2005), soil inocula for each level of phylogenetic distance were necessarily sourced from different species.

**Figure 1.**
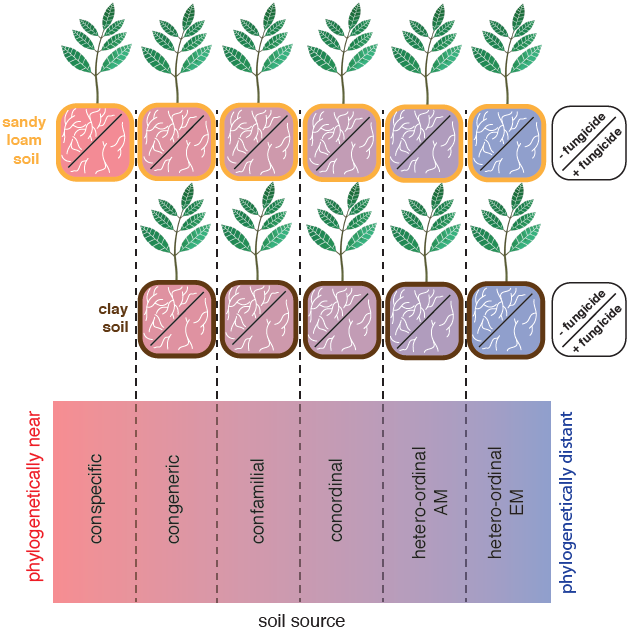
Experimental design overview. Schematic diagram illustrating the different experimental treatments used in the study. For each of the eight species used in the experiment, the figures shows the factorial arrangement of soil type (sandy loam, clay), fungicide treatments (+/-) and phylogenetic distance to the overstory tree from beneath which soils were sourced, ranging from conspecific (pink) to hetero-ordinal (purple).

We took two approaches to separate the effects of the soil microbiota and nutrient availability on variation in PSF. First, as all mycorrhizas and many key plant pathogens are fungal, we added a factorial fungicide treatment such that half of all seedlings of each species in each soil type × phylogenetic distance treatment combination received fungicide. Second, for each soil source, we carefully measured starting concentrations of several soil nutrients. In total, our experiment consisted of ∼1200 seedlings, with an average of 3-10 seedlings per treatment combination, depending on species (Table S1).

### Seedling experiment

In February and March of 2014, seedlings were transplanted from germination trays into soil mixtures in 2-L polyethylene pots in the shadehouse. Transplanting into soil mixtures took place within three days of inoculum collection. Within each treatment combination, seedlings were distributed evenly among 16 nursery benches. Whole benches were designated either for control (non-fungicide) or fungicide treatment, alternating control and fungicide benches inside the shadehouse, to avoid fungicide drift during application. Pairs of adjacent control and fungicide benches were designated as an experimental block, for a total of eight experiment blocks. From ten days after transplanting to the end of the experiment, half of all seedlings of a species across treatment combinations were treated with CAPTAN fungicide every two weeks. Fungicide was applied as a soil drench in aliquots of 50ml / individual at a rate of 0.66g/L following (Liu *et al.* 2011). All control plants received a 50ml “mock treatment” of water at the same interval. We quantified the supply rates of 14 anions and cations (see Fig S1) in each soil type × fungicide treatment combination using Plant Root Simulator Probes (PRS), which use ion exchange resin membranes (Western Ag Innovations Inc., Saskatoon, SK, Canada). Probes were buried in 2L pots without seedlings for 8 weeks.

### Seedling harvest

Experimental seedlings were harvested between mid-February and June 2015, and ranged from 325-453 days old (Appendix S1). We oven dried all tissues at 60° C for at least 72 hours before weighing to an accuracy of 0.0001g to obtain the biomasses of leaves, stems, and coarse and fine roots and total plant biomass. To develop allometric equations for estimation of initial biomass of experimental seedlings, the same data were collected for a subset of 6-20 seedlings (depending on seed availability) of each, which were harvested at the initial transplant (Appendix S1).

Fresh roots of all EM host species (*Dryobalanops, S. curtisii, S. laxa, S. scaberrima, Castanopsis*) were inspected for ECM colonization at harvest using a 10X-dissecting microscope. We estimated percent colonization by counting approximately 100 root tips from 10 *ca.* 1-cm root fragments sampled haphazardly from the root system of each seedling, and scoring each tip as either colonized or uncolonized. Estimates of colonization were made immediately following root washing, and root fragments used were returned to the root system before further processing. Field conditions prevented us from estimating AM colonization, which requires additional staining procedures and higher magnification microscopy.

### Data Analysis – Seedling-soil phylogenetic distances

We estimated evolutionary relationships among tree species in this study by assembling a phylogenetic tree of all plant species used either as focal species or soil inocula using a tree pruned from (Zanne et al. 2014) (Appendix S1). We calculated phylogenetic distance between all pairs of species using the package “ape” (Paradis *et al.* 2004) and used this pairwise distance matrix to estimate the divergence time between each focal seedling and the species from which we had sourced the inoculum in which it had been grown. Seedling-soil phylogenetic distances were standardized relative to the maximum of the seedling-soil phylogenetic distances, expressed as divergence times, across all focal species (361 mya) to give a relative phylogenetic distance measurement ranging from 0 (conspecific) to 1 (distantly related) (Liu et al. 2011).

### Soil fertility

Since soil nutrient rates were correlated (Fig S1), we used the first component of a principle component analysis as an index of soil fertility as a predictor in analyses. The first principle component explained 78.4% of variance in soil nutrient availability, and was positively correlated with nitrate, phosphorous, potassium, calcium, and magnesium (Figure S2). We used Moran’s I to confirm that there was no phylogenetic signal in soil fertility (Appendix S1).

### Growth analysis

We calculated relative growth rate of biomass (mg g^-1^ day^-1^), hereafter RGR, as [log(*bm*_*h*_) – log (*bm*_*i*_)] *d*^-1^, where *bm*_*h*_ is biomass at harvest, *bm*_*i*_ is estimated initial biomass at first census (at transplant), and *d* is the number of days between the initial census and harvest. Initial biomass estimates were predicted from species-specific linear regression models developed using the *stem diameter, stem height*, and *leaf number* measured on seedlings harvested at transplant as predictor variables. We selected reduced models for each species using an AIC threshold of 2 and used the resulting preferred models to estimate initial biomass for seedlings in the experiment (adjusted *R*^*2*^ = 0.32-0.97, Table S2).

To measure treatment effects on seedling growth rate we used linear mixed-effects models with RGR of total biomass as a response and mycorrhizal type (AM vs EM), seedling-soil relative phylogenetic distance, soil fertility index, and fungicide treatment (control/no-fungicide vs. fungicide) as fixed effects. We retained all higher order interactions among fixed effects, as they were explicitly designed into the experiment to test our hypotheses. Species and experimental block were modeled as random effects with a nested structure (block within species). Support for different random effects models was evaluated using AIC model selection with maximum likelihood estimation, which favored retaining random intercepts and dropping random slopes for both variables. To better evaluate individual species responses to experimental treatments, we also fit separate mixed models for each focal species the same fixed effect model, but with only experimental block as a random intercept.

All growth models were fit using the package *nlme* (Pinheiro *et al.* 2018). Residuals were assessed visually for normality and heteroscedasticity and using Levene’s tests for equality of variances. Where necessary, heteroscedasticity of continuous or within-group error was modeled using variance functions in the *nlme* package, and support for variance weighting was confirmed using AIC. We compared fixed effects estimates across experimental groups using least squares means to estimate the marginal effect of linear predictors as implemented in the package *emmeans* (Lenth 2016).

### EM colonization analysis

We analyzed effects of all experimental treatments on percent EM colonization of all focal species using a GLMM of the number of colonized root tips on a seedling out of 100 sampled. We used the same model structure as for the growth models described above, except with a negative binomial error distribution. We used a negative binomial because a binomial error distribution was not a good fit to the percent colonization data due to overdispersion, as supported by AIC model selection and the dispersion parameter estimate (Table S4). To correct for zero inflation, we dropped zero observations for one species (*Castanopsis*) and separately modeled colonization success (whether a seedling was colonized or not) for *Castanopsis* using a generalized linear mixed model (GLMM) with a binomial error distribution (Appendix S1).

To assess the effects of EM colonization on seedling RGR, we used simple linear regression model with EM host species and percent root colonization as interacting fixed effects. All GLMMs were fit using the packages “lme4” (Bates *et al.* 2015), and dispersion parameters were estimated using the package “blmeco” (Korner-Nievergelt *et al.* 2015). We compared marginal effects estimates of fixed factors across experimental groups as implemented in the packages “emmeans” (Lenth 2016) and “sjPlot” (Lüdecke 2018).

## Results

### Growth response

Seedling-soil phylogenetic distance affected seedling RGR differently in AM versus EM groups, and this difference was removed by fungicide application (significant mycorrhizal type × phylogenetic distance × fungicide interaction; Table 1, Figure 2). In EM control seedlings, RGR decreased as the phylogenetic distance increased, consistent with positive PSF (phylogenetic distance slope = -0.63 ± 0.20). In contrast, for AM seedlings, biomass growth rate increased as phylogenetic distance increased, consistent with negative PSF (phylogenetic distance slope = 0.51 ± 0.48). Among fungicide-treated seedlings, phylogenetic distance did not significantly affect the biomass growth rate of seedlings of either mycorrhizal type (EM phylogenetic distance slope = - 0.16 ± 0.29, AM phylogenetic distance slope = -0.24 ± 0.33). The impact of phylogenetic distance also varied with soil fertility, depending on mycorrhizal type (significant mycorrhizal type × soil fertility × phylogenetic distance interaction; Table 1, Figure 3). While variation in biomass growth of AM seedlings with phylogenetic distance tended to be more consistent with negative PSF as soil fertility increased, for EM seedlings the growth-phylogenetic distance relationship did not depend on soil fertility.

**Table 1:** Treatment effects on seedling biomass growth rate based on linear mixed model ANCOVA using fixed effects of host *mycorrhizal type* (AM or EM), *soil fertility*, seedling-soil *phylogenetic distance*, and *fungicide* application with random effects of *focal species* and *experimental block*.

| Fixed effects | $X^2$ | df | $p$ |
| --- | --- | --- | --- |
| Mycorrhizal Type | 0.137 | 1 | 0.711 |
| Soil Fertility | 0.792 | 1 | 0.374 |
| Phylogenetic Distance | 4.591 | 1 | 0.032 |
| Fungicide | 1.113 | 1 | 0.291 |
| Mycorrhizal type : Soil fertility | 3.692 | 1 | 0.055 |
| Mycorrhizal type : Phylogenetic distance | 3.028 | 1 | 0.082 |
| Soil fertility : Phylogenetic distance | 2.682 | 1 | 0.101 |
| Mycorrhizal type : Fungicide | 0.403 | 1 | 0.525 |
| Soil fertility : Fungicide | 0.489 | 1 | 0.484 |
| Phylogenetic distance : Fungicide | 0.019 | 1 | 0.889 |
| Mycorrhizal type : Soil fertility : Phylogenetic distance | 4.502 | 1 | 0.034 |
| Mycorrhizal type : Soil fertility : Fungicide | 1.380 | 1 | 0.240 |
| Mycorrhizal type : Phylogenetic distance : Fungicide | 4.180 | 1 | 0.041 |
| Soil fertility : Phylogenetic distance : Fungicide | 1.089 | 1 | 0.297 |
| Mycorrhizal type : Soil fertility : Phylogenetic distance : Fungicide | 0.034 | 1 | 0.853 |
| Random effects | Variance | SD |  |
| Focal species | 1.479 | 1.216 |  |
| Block | 0.228 | 0.48 |  |

**Figure 2.**
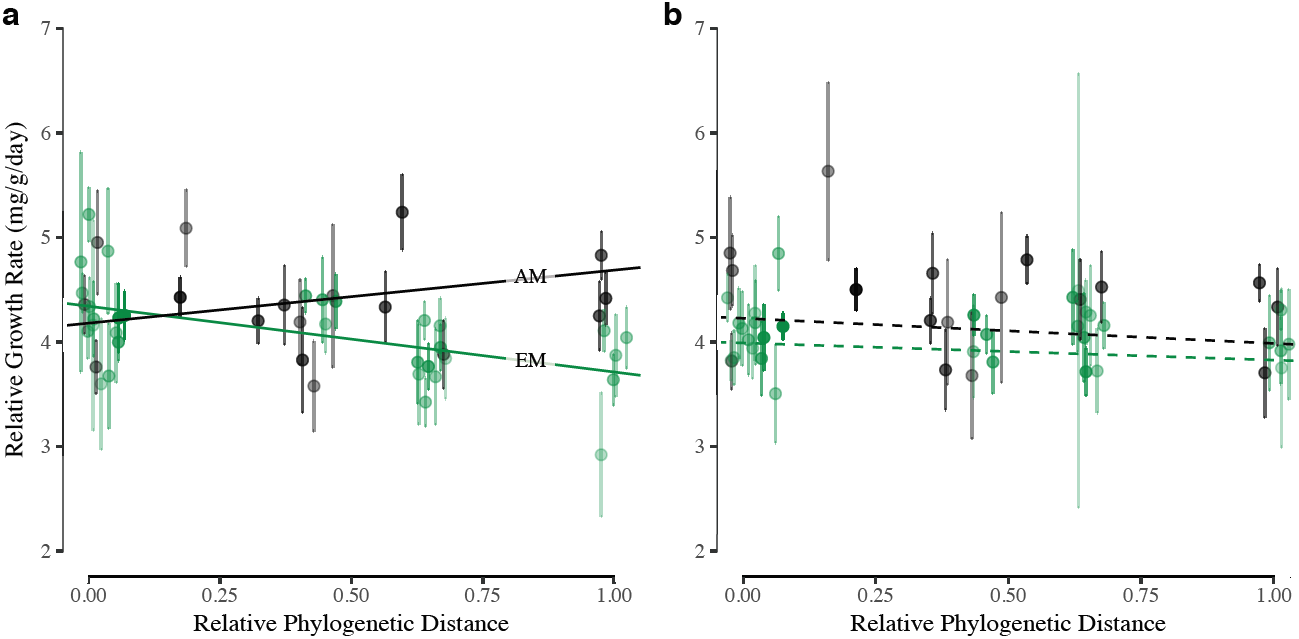
Plant-soil phylogenetic distance effect on plant growth across mycorrhizal types in control (a) and fungicide (b) groups. AM group is shown in black, EM in green. Data are summarized as the within species by phylogenetic distance mean ± SE to improve readability and include the random effects; point intensity indicates the number of observations within each summary point (range= 3-27). Trend lines indicate the estimated marginal effect of seedling-soil phylogenetic distance for treatment combinations; solid lines indicate estimates not overlapping zero, while dashed lines indicate estimates overlapping zero. *Mycorrhizal type* × *phylogenetic distance* × *fungicide* treatment interaction p=0.041. Trend lines estimate marginal effects at intermediate fertility.

**Figure 3.**
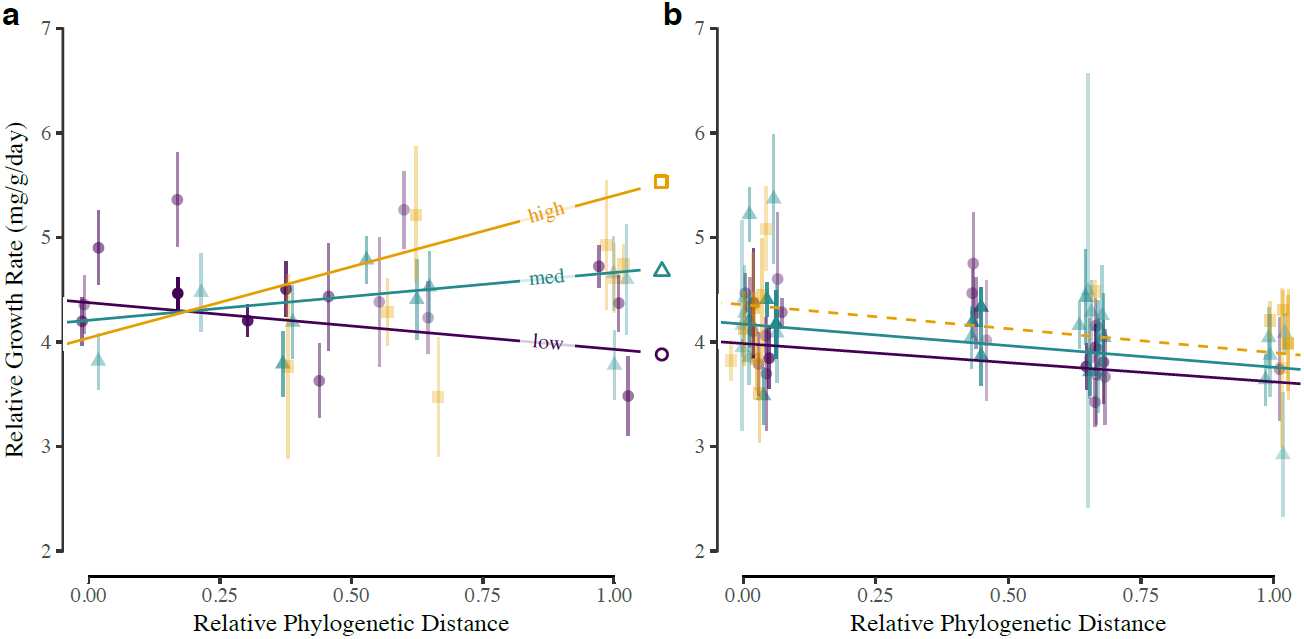
Plant-soil phylogenetic distance effect on plant growth across at different soil fertility levels in AM (panel a) and EM (panel b) groups. Purple circle, blue triangle, and yellow squares indicate low, intermediate, and high soil fertility. Data are summarized as the within species X phylogenetic distance mean +- SE, binned over three soil fertility intervals to improve readability and have been adjusted for random effects; transparency indicates observations within each summary point (range= 3-37). Trend lines indicate the estimated marginal effect of seedling-soil phylogenetic distance; solid lines indicate estimates not overlapping zero, while dashed lines indicate estimates overlapping zero. *Mycorrhizal type* ×*soil fertility* × *phylogenetic distance* interaction p=0.034. Data and trend lines were averaged over both fungicide and control groups.

In the all-species model of biomass growth, the random effects model incorporating species and experimental block accounted for most of the explained variation in growth rate, whereas fixed effects accounted for only 2% (*marginal-R*^*2*^of 54% vs. *conditional-R*^*2*^ of 56%). For individual species models, effect sizes for experimental treatments differed greatly among species (Table S5), and the variance explained by fixed effects ranged from negligible, *marginal-R*^*2*^ < 0.1% (*Castanopsis*) to *marginal-R*^*2*^ = 14% (*S. scaberrima*), with seedling-soil phylogenetic distance accounting for < 0.1% (*Madhuca*) to 10% (*S. scaberrima*). In two species, second order interactions explained more variance than all main treatment effects combined (*Dacryodes, Dryobalanops*).

### Ectomycorrhizal Colonization

We found that in EM seedlings, all three main effects of soil fertility, phylogenetic distance, and fungicide application significantly affected the percent EM colonization of seedling root systems (Table S6). Fungicide application significantly reduced colonization by EM fungi (*X*^2^=19.29, df=1, p<0.001) by roughly six percent (39.5 ± 1.6 to 33.3 ± 1.4) (Fig 4c), suggesting that fungicide treatment had direct effects on root-associated fungi. Increasing seedling-soil phylogenetic distance significantly reduced root fungal colonization in EM hosts (*X*^*2*^ = 9.37, df = 1, p=0.002) (Fig 4a). Increased soil fertility was associated with greater colonization (*X*^*2*^=18.62, df=1, p<0.001), and this effect was more pronounced in control seedlings than those that received fungicide (interaction *X*^*2*^=10.12, df=1, p=0.001)(Fig 4b). Colonization was also significantly associated with faster biomass growth rate (F_1,599_=55.27, p<0.001), an effect that was consistent across species (interaction F_4,599_=0.5325, p=0.71) (Fig 5). In species-specific tests, the correlation between root colonization and growth varied substantially and was significant for all species except *Castanopsis* (*Castanopsis* r=0.17, *df* = 59, p=0.186, *Dryobalanops* r=0.25, *df* = 145, p=0.002, *S. curtisii* r(187)=0.47, p<0.001, *S. laxa* r(143)=0.20, p=0.014, *S. scaberrima* r(65)=0.27, p=0.030).

**Figure 4:**
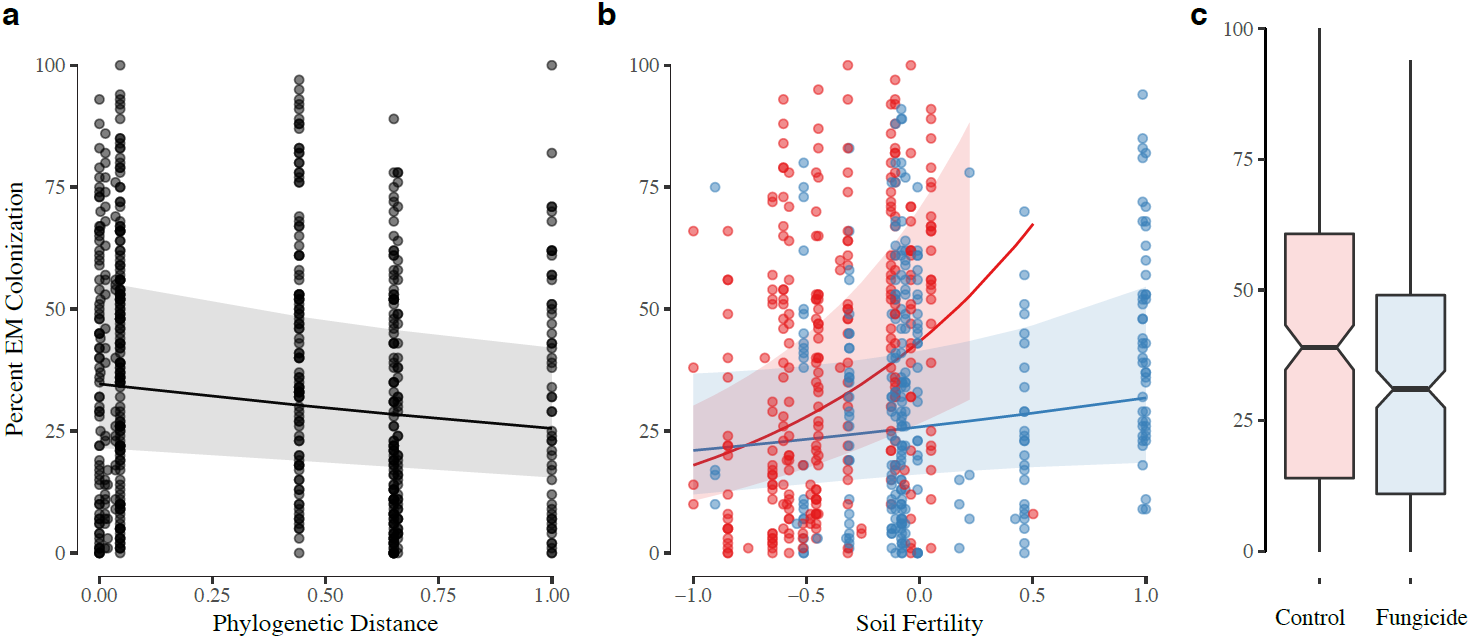
Treatment effects on root colonization by EMF. In panels **b** and **c**, control treatments are represented in red, while fungicide treatments are represented in blue. Trend lines in panels **a** and **b** indicate generalized mixed model predictions of root colonization and shaded regions indicate 95% confidence intervals.

**Figure 5:**
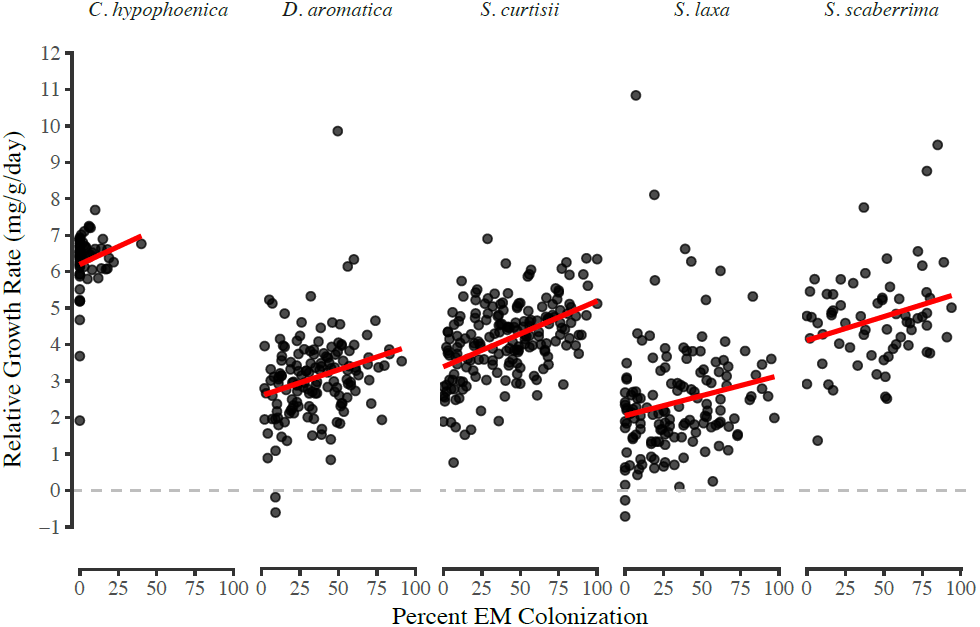
Root colonization by EMF is positively associated with seedling growth rate across all EM species. Fit lines indicate linear model fits for each species.

## Discussion

Negative plant soil feedbacks promote coexistence in high diversity plant communities owing to the strong effects of species-specific fungal pathogens (Augspurger & Wilkinson 2007; Bagchi et al. 2010b; 2014). Despite recognition of the effects of EMF on both PSF (Bennett et al. 2017) and critical soil processes (Clemmensen et al. 2013; Phillips et al. 2013; Averill et al. 2014; Terrer et al. 2016), there are few studies of tropical PSF outside of AM-dominated neotropical forests. To address these knowledge gaps, we used a manipulative experiment with seedlings of eight tree species to investigate sources of variation in PSFs in a high diversity, co-dominant AM-EM rain forest in Borneo. Our results show that seedling recruitment in this forest is influenced by PSFs. Moreover, we found that the strength and direction of PSFs depend strongly on both tree species’ mycorrhizal association and the local soil environment, defined by nutrient availability and the phylogenetic relatedness of the overstory tree species.

The strength of CNDD and PSF experienced by tropical tree species is often interpreted as primarily a function of susceptibility to natural enemies. However, we show that PSFs are far more complicated than this. Considering seeds dispersed from a mother tree in a natural forest, we propose that the landscape of spatial variation in PSF that they encounter on the forest floor and that affects seedling performance is shaped by diverse sources of variation. Many of these sources of variation stem not only from negative interactions with natural enemies, but also, as our results imply, from how soil resource availability and evolutionary history affect the nature of resource-trading interactions between plants and fungal mutualists, which also depend on the plant’s endogenous resource-allocation strategy and fungal physiology. Theoretical models show that the size of the tree species pool combined with differences in the benefits conferred by AM vs EM fungi to their hosts affect the probability of tree species’ monodominance (Fukami et al. 2017). Yet, how the complexities that we have discovered in the drivers of PSF enhance the diversity-maintaining mechanism of CNDD versus promote dominance of tree taxa with particular mycorrhizal associations across environmentally heterogeneous landscapes requires further exploration.

The regional dominance of species in a single tree family, the Dipterocarpaceae, in southeast Asian forests (Alexander & Hogberg 1986), as well as the capacity of some EM host species to form monodominant stands in otherwise high-diversity tropical rain forests (Connell & Lowman 1989; Torti et al. 2001) have defied convincing explanation. One hypothesis is that, in contrast to AM host species, EM host species have less negative or even positive PSFs, and that EM PSFs become more positive in soils conditioned by closely related species, which would ultimately act to generate a positive feedback for recruitment of EM host species at larger spatial scales (Fukami *et al.* 2017). Consistent with this hypothesis, we demonstrate that seedlings of EM species, particularly those of the Dipterocarpaceae, experience more positive PSFs and that association with fungal mutualists likely underlies these effects. We also found that seedlings of AM host species experienced more negative PSF and that the effect of phylogenetic distance for them was opposite to EM host species. Fundamentally, the suite of possible drivers of PSFs must be the same across all forests, but we have demonstrated here, for the first time, that the relative strength, direction, and phylogenetic signal in PSF differs across mycorrhizal types and soil environments in a species-rich tropical forest. By extending the generality of findings from temperate forests that show significant variation in PSFs across mycorrhizal types (Bennett et al. 2017), our results suggest that there are important biogeographic differences across major tropical regions in the relative importance of different drivers of PSFs and how they ultimately shape forest community structure.

### Variation in feedback effects

There is increasing evidence that the abundance and distribution of EM host species can have profound impacts on ecosystem function and patterns of diversity in forests (Averill *et al.* 2014; Bennett *et al.* 2017). Previous work in temperate North America has indicated that EM and AM tree species may experience different plant-soil dynamics, with EM and AM taxa experiencing conspecific facilitation and inhibition, respectively (Bennett et al. 2017). Given the substantial differences climate, environment, and biotic assemblages, it was unclear whether similar patterns would also be observed in tropical ecosystems, but our findings are consistent with those from temperate forests and extend them by showing variation between AM and EM host species in the effect of phylogenetic relatedness on PSF. For seedlings of EM trees, slower growth in soil inocula from under distantly related taxa likely results from reduced availability or compatibility of EMF inoculum in soil and thus reduced root colonization by these mutualists, as we showed. While previous work in this forest has shown that interactions between EM fungi and Dipterocarpaceae may not show strict host specificity at the species-level (Peay et al. 2015), our results here suggest that host preferences exist that can improve colonization and benefits for seedlings growing beneath conspecifics or close relatives, leading to positive PSFs. The beneficial effects we observed could also stem from reduced pathogen infection, as EM fungi can increase disease resistance, including via direct antagonism with soil fungal pathogens (Marx 1972; Chakravarty & Unestam 1987; ZengPu & JunRan 1995). Thus, while all seedlings likely confront greater abundance of pathogens to which they are susceptible when growing near close relatives, our results suggest that such negative effects could be more strongly ameliorated in EM versus AM hosts by improved growth resulting from greater EMF colonization or more favorable resource-trading relationships and reduced infection by or impacts of pathogens.

In contrast, we show the opposite pattern in AM hosts, of faster growth in soil inocula from under distantly related taxa, which could arise through several mechanisms. Since closely related plant species tend to share pathogens, this phylogenetic PSF could be mediated by escape from natural enemies of near relatives (Gilbert & Webb 2007; Liu et al. 2011). Since AM fungi are generally broader in host-range than EM fungi (Davison et al. 2015), seedlings of AM host species are less likely to lose the benefit of mutualists when growing under more distantly related tree species. Indeed, because of variation along the mutualism-parasitism continuum (Bronstein 1994), AM fungi conferring minimal benefits to plants can dominate locally and cause negative feedbacks (Bever 2002). Dissecting how the interactions with pathogens and mutualists differ between AM and EM host species to produce the variation in PSFs that we observed would be aided by genomic evidence identifying the microbial colonists of these seedlings’ roots.

Fungicide application negated both the phylogenetic PSF observed in control seedlings, and the difference in PSF observed between EM and AM host species, consistent with a fungal origin of these effects. This was also supported by the reduced EM colonization of roots, suggesting that fungicide application reduced EM, and likely AM, fungal abundance in soils. Although fungi were not totally excluded, fungicides are known to vary tremendously in their efficiency (e.g. (Bagchi *et al.* 2014).

Individual species were also a significant source of variation in PSF strength. This is not surprising, given the individual variation in growth between species, and that variation in PSF within mycorrhizal types has been documented in the temperate biome as well (Bennett et al. 2016). This interspecific variability has ecological consequences, but also highlights the importance of testing multiple species in PSF experiments given that major effects were not detectable with the reduced power of some single species models.

### Interactive effects of biotic feedback and resource availability

Ecological theory and empirical evidence indicate that resource availability can strongly influence species interactions and may shift crucial balance points for interactions that exist along a mutualism-parasitism continuum, including those between plants and mycorrhizal fungi (Johnson *et al.* 1997). Soil fertility may affect CNDD, with stronger CNDD associated with greater soil resource availability in a temperate forest (LaManna et al. 2016). Consistent with this, we found that among AM host species, greater soil resource availability was associated with more negative PSFs. Johnson et al. (2010) also found that AMF of temperate grass species became less beneficial as soil fertility increased. Among EM species, however, we found that soil fertility had no effect on PSF. While the mechanisms controlling resource trading relationships between plants and EM fungi are poorly understood, our finding is consistent with the view that, compared to AM, EM associations are more beneficial to plants (Bennett et al. 2016; but see (Karst et al. 2008)) and with greater plant control (Nehls et al. 2007). Alternatively, EMF may simply have different functionality that provides similar benefits across the range of soil conditions in our study. In support of this idea, a recent study showed that tropical EM-associated seedlings assimilated P in a broader range of chemical forms than AM-associated seedlings (Liu *et al.* 2018). All soils in our study site are relatively nutrient depleted, so it is possible that different outcomes would have been observed at the high nitrogen levels that have been shown in temperate forests to disrupt EM symbiosis (van der Linde et al. 2018).

### Biogeographic variation in PSF

Our findings of positive phylogenetic PSFs may help explain some of the biogeographic differences among tropical rainforest. Average CNDD has been observed to be strongest in species rich communities and at lower latitudes (Johnson *et al.* 2012), an effect which is often attributed to more negative effects of pathogenic fungi nearer the equator (Bagchi et al. 2014, Bennett et al. 2016). In neotropical forests, where the Janzen-Connell hypothesis was first developed, negative PSFs appear to predominate (Mangan et al. 2010). While we observed some negative PSFs for AM host species, PSFs varied strongly by mycorrhizal type and soil fertility, raising questions about the generality of negative PSFs and whether they are the dominant process maintaining species diversity in tropical tree communities. While the monospecific dominance of EM hosts observed in some neotropical forests is associated with a positive home soil advantage (McGuire et al. 2007; Corrales et al. 2016), we show that positive phylogenetic PSF exist in a tropical forest in which a high diversity of both EM and AM host species coexist. This positive phylogenetic feedback may explain some of the unique features of SE Asian forests, such as the familial dominance of the Dipterocarpaceae. Similalry, tree species distributions in dipterocarp forests are more spatially aggregated than in non-dipterocarp forests (Condit et al. 2000), which could arise from limited dispersal, but only in combination with more positive PSFs for EM host species, as we found. However, it is unclear why some systems exhibit monospecific dominance of EM host species, whereas others exhibit co-existence of AM and EM species at comparable levels of richness. Fukami et al. (2017) developed a non-phylogenetic PSF model showing that such variation can arise due to differences in trait characteristics of regional species pools that would be promising to follow up with future empirical studies.

## Conclusions

In this study, we have made explicit consideration of fungal mutualism and soil fertility in mediating PSFs in a high diversity tropical rainforest. We demonstrate that host mycorrhizal association interacts with soil fertility to produce variable phylogenetic PSFs. To the extent that PSF contribute to CNDD, they must influence diversity in plant communities (Mills & Bever 1998; Johnson *et al.* 2012), and negative PSF are often viewed as a primary driver of CNDD in species-rich tropical forests (Mangan *et al.* 2010b). Our documentation of positive and highly variable PSF in a diverse tropical forest helps explain biogeographic differences in forest structure across tropical rainforests, but also calls for greater investigation into the generality of negative PSF in maintaining tropical tree diversity.

## Acknowledgements

This research was funded by NSF RAPID grant 1361171 to KGP and SER. The authors would like to thank the Sarawak Forest Research Corporation and Forest Department Sarawak for permission to conduct research at Lambir Hills. We also thank park warden Januarie Kulis as well as the entire staff of Lambir Hills National Park, Lip Khoon Kho of the Tropical Peat Research Institute, and Lilyen Ukat for their help and logistical support of this research. We extend our deepest gratitude to the community of Sungai Liam for their gracious hospitality and logistical support of this work. We also thank members of the Peay Lab group for helpful review and feedback on early drafts of this manuscript.

## References

1. Alexander, I.J. & Hogberg, P. (1986). Ectomycorrhizas of tropical angiospermous trees. New Phytologist, 541–549.

2. Augspurger, C.K. (1983). Seed Dispersal of the Tropical Tree, Platypodium Elegans, and the Escape of its Seedlings from Fungal Pathogens. Journal of Ecology, 71, 759–771.

3. Augspurger, C.K. & Wilkinson, H.T. (2007). Host Specificity of Pathogenic Pythium Species: Implications for Tree Species Diversity. Biotropica, 39, 702–708.

4. Averill, C., Turner, B.L. & Finzi, A.C. (2014). Mycorrhiza-mediated competition between plants and decomposers drives soil carbon storage. Nature, 505, 543–545.

5. Bagchi, R., Gallery, R.E., Gripenberg, S., Gurr, S.J., Narayan, L., Addis, C.E., et al. (2014). Pathogens and insect herbivores drive rainforest plant diversity and composition. Nature, 506, 85–88.

6. Bagchi, R., Press, M.C. & Scholes, J.D. (2010a). Evolutionary history and distance dependence control survival of dipterocarp seedlings. Ecology Letters, 13, 51–59.

7. Bagchi, R., Swinfield, T., Gallery, R.E., Lewis, O.T., Gripenberg, S., Narayan, L., et al. (2010b). Testing the Janzen-Connell mechanism: pathogens cause overcompensating density dependence in a tropical tree. Ecology Letters, 13, 1262–1269.

8. Baillie, I.C., Ashton, P.S., Chin, S.P., Davies, S.J., Palmiotto, P.A., Russo, S.E., et al. (2006). Spatial associations of humus, nutrients and soils in mixed dipterocarp forest at Lambir, Sarawak, Malaysian Borneo. J. Trop. Ecol., 22, 543–553.

9. Bates, D., Mächler, M., Bolker, B. & Walker, S. (2015). Fitting Linear Mixed-Effects Models Using lme4. J. Stat. Soft., 67, 1–48.

10. Bennett, J.A., Maherali, H., Reinhart, K.O., Lekberg, Y., Hart, M.M. & Klironomos, J.N. (2017). Plant-soil feedbacks and mycorrhizal type influence temperate forest population dynamics. Science, 355, 181–184.

11. Bever, J.D. (2002). Negative feedback within a mutualism: host-specific growth of mycorrhizal fungi reduces plant benefit. Proceedings of the Royal Society B: Biological Sciences, 269, 2595–2601.

12. Bever, J.D., Westover, K.M. & Antonovics, J. (1997). Incorporating the Soil Community into Plant Population Dynamics: The Utility of the Feedback Approach. The Journal of Ecology, 85, 561.

13. Brearley, F.Q. (2012). Ectomycorrhizal Associations of the Dipterocarpaceae. Biotropica, 44, 637–648.

14. Bronstein, J.L. (1994). Conditional outcomes in mutualistic interactions. Trends in Ecology & Evolution, 9, 214–217.

15. Bruns, T.D. & Shefferson, R.P. (2004). Evolutionary studies of ectomycorrhizal fungi: recent advances and future directions. Can. J. Bot., 82, 1122–1132.

16. Bruns, T.D., Bidartondo, M.I. & Taylor, D.L. (2002). Host specificity in ectomycorrhizal communities: what do the exceptions tell us? Integr. Comp. Biol., 42, 352–359.

17. Chakravarty, P. & Unestam, T. (1987). Differential influence of ectomycorrhizae on plant growth and disease resistance in Pinus sylvestris seedlings. Journal of Phytopathology.

18. Clemmensen, K.E., Bahr, A., Ovaskainen, O., Dahlberg, A., Ekblad, A., Wallander, H., et al. (2013). Roots and Associated Fungi Drive Long-Term Carbon Sequestration in Boreal Forest. Science, 339, 1615–1618.

19. Comita, L.S., Muller-Landau, H.C., Aguilar, S. & Hubbell, S.P. (2010). Asymmetric Density Dependence Shapes Species Abundances in a Tropical Tree Community. Science, 329, 330–332.

20. Condit, R., Ashton, P.S., Baker, P., Bunyavejchewin, S., Gunatilleke, S., Gunatilleke, N., et al. (2000). Spatial Patterns in the Distribution of Tropical Tree Species. Science, 288, 1414–1418.

21. Condit, R., Hubbell, S.P. & Foster, R.B. (1992). Recruitment Near Conspecific Adults and the Maintenance of Tree and Shrub Diversity in a Neotropical Forest. Am Nat, 140, 261–286.

22. Connell, J.H. (1971). On the role of natural enemies in preventing competitive exclusion in some marine animals and in rain forest trees. Dynamics of populations, 298, 312.

23. Connell, J.H. & Lowman, M.D. (1989). Low-diversity tropical rain forests: some possible mechanisms for their existence. Am Nat.

24. Davies, S.J., Tan, S., LaFrankie, J.V. & Potts, M. (2005). Soil-related floristic variation in a hyperdiverse dipterocarp forest. In: Pollination ecology and rain forest diversity, Sarawak studies. Springer, New York, pp. 22–34.

25. Davison, J., Moora, M., Öpik, M., Adholeya, A., Ainsaar, L., Bâ, A., et al. (2015). Global assessment of arbuscular mycorrhizal fungus diversity reveals very low endemism. Science, 349, 970–973.

26. Fukami, T., Nakajima, M., Fortunel, C., Fine, P.V.A., Baraloto, C., Russo, S.E., et al. (2017). Geographical Variation in Community Divergence: Insights from Tropical Forest Monodominance by Ectomycorrhizal Trees. Am Nat, 190, S105–S122.

27. Gilbert, G.S. & Webb, C.O. (2007). Phylogenetic signal in plant pathogen-host range. Proc. Natl. Acad. Sci. U.S.A., 104, 4979–4983.

28. Gilbert, G.S., Briggs, H.M. & Magarey, R. (2015). The Impact of Plant Enemies Shows a Phylogenetic Signal. PLoS ONE, 10, e0123758–11.

29. Hoeksema, J.D., Chaudhary, V.B., Gehring, C.A., Johnson, N.C., Karst, J., Koide, R.T., et al. (2010). A meta-analysis of context-dependency in plant response to inoculation with mycorrhizal fungi. Ecology Letters, 13, 394–407.

30. Ishida, T.A., Nara, K. & Hogetsu, T. (2007). Host effects on ectomycorrhizal fungal communities: insight from eight host species in mixed conifer–broadleaf forests. New Phytol., 174, 430–440.

31. Janos, D.P. (1980). Mycorrhizae influence tropical succession. Biotropica, 12, 56.

32. Janzen, D.H. (1970). Herbivores and the number of tree species in tropical forests. Am Nat, 501–528.

33. Johnson, D.J., Beaulieu, W.T., Bever, J.D. & Clay, K. (2012). Conspecific negative density dependence and forest diversity. Science.

34. Johnson, N.C., Graham, J.H. & Smith, F.A. (1997). Functioning of mycorrhizal associations along the mutualism-parasitism continuum. New Phytologist, 135, 575–586.

35. Johnson, N.C., Wilson, G.W.T., Bowker, M.A., Wilson, J.A. & Miller, R.M. (2010). Resource limitation is a driver of local adaptation in mycorrhizal symbioses. Proc. Natl. Acad. Sci. U.S.A., 107, 2093–2098.

36. Karst, J., Marczak, L., Jones, M.D. & Turkington, R. (2008). The mutualism-parasitism continuum in ectomycorrhizas: a quantitative assessment using meta-analysis. Ecology, 89, 1032–1042.

37. Klironomos, J.N. (2002). Feedback with soil biota contributes to plant rarity and invasiveness in communities. Nature, 417, 67–70.

38. Korner-Nievergelt, F., Roth, T., Felten, Von, S. & Guélat, J. (2015). Bayesian data analysis in ecology using linear models with R, BUGS, and Stan.

39. LaManna, J.A., Walton, M.L., Turner, B.L. & Myers, J.A. (2016). Negative density dependence is stronger in resource-rich environments and diversifies communities when stronger for common but not rare species. Ecology Letters, 1–11.

40. Lee, H.S., Davies, S.J., LaFrankie, J.V., Tan, S. & Yamakura, T. (2002). Floristic and structural diversity of mixed dipterocarp forest in Lambir Hills National Park, Sarawak, Malaysia. Journal of Tropical Forest ….

41. Lenth, R.V. (2016). Least-Squares Means: The RPackage lsmeans. J. Stat. Soft., 69, 1–33.

42. Liu, X., Burslem, D.F.R.P., Taylor, J.D., Taylor, A.F.S., Khoo, E., Lee, N.M., et al. (2018). Partitioning of soil phosphorus among arbuscular and ectomycorrhizal trees in tropical and subtropical forests. Ecology Letters, 21, 713–723.

43. Liu, X., Liang, M., Etienne, R.S., Wang, Y., Staehelin, C. & Yu, S. (2011). Experimental evidence for a phylogenetic Janzen-Connell effect in a subtropical forest. Ecology Letters, 15, 111–118.

44. Liu, Y., Fang, S., Chesson, P. & He, F. (2015). The effect of soil-borne pathogens depends on the abundance of host tree species. Nat Commun, 6, 10017.

45. Lüdecke, D. (2018). sjPlot: Data Visualization for Statistics in Social Science. R package version 3.1–137.

46. Mangan, S.A., Herre, E.A. & Bever, J.D. (2010a). Specificity between Neotropical tree seedlings and their fungal mutualists leads to plant-soil feedback. Ecology, 91, 2594–2603.

47. Mangan, S.A., Schnitzer, S.A., Herre, E.A., Mack, K.M.L., Valencia, M.C., Sanchez, E.I., et al. (2010b). Negative plant–soil feedback predicts tree-species relative abundance in a tropical forest. Nature, 466, 752–755.

48. Marx, D.H. (1972). Ectomycorrhizae as biological deterrents to pathogenic root infections. Annu Rev Phytopathol, 10, 429–454.

49. McGuire, K.L. (2007). Common ectomycorrhizal networks may maintain monodominance in a tropical rain forest. Ecology, 88, 567–574.

50. Mehrabi, Z. & Tuck, S.L. (2014). Relatedness is a poor predictor of negative plant-soil feedbacks. New Phytologist, /a–n/a.

51. Metz, M.R., Sousa, W.P. & Valencia, R. (2010). Widespread density-dependent seedling mortality promotes species coexistence in a highly diverse Amazonian rain forest. Ecology, 91, 3675–3685.

52. Mills, K.E. & Bever, J.D. (1998). Maintenance of diversity within plant communities: soil pathogens as agents of negative feedback. Ecology, 1595–1601.

53. Morris, W.F., Hufbauer, R.A., Agrawal, A.A., Bever, J.D., Borowicz, V.A., Gilbert, G.S., et al. (2007). Direct and interactive effects of enemies and mutualists on plant performance: a meta-analysis. Ecology, 88, 1021–1029.

54. Moyersoen, B., Becker, P. & Alexander, I.J. (2001). Are ectomycorrhizas more abundant than arbuscular mycorrhizas in tropical heath forests? New Phytologist.

55. Nara, K. (2006). Ectomycorrhizal networks and seedling establishment during early primary succession. New Phytol., 169, 169–178.

56. Newbery, D.M., Alexander, I.J., Thomas, D.W. & Gartlan, J.S. (1988). Ectomycorrhizal rain-forest legumes and soil phosphorus in Korup National Park, Cameroon. New Phytol., 109, 433–450.

57. Paradis, E., Claude, J. & Strimmer, K. (2004). APE: Analyses of Phylogenetics and Evolution in R language. Bioinformatics, 20, 289–290.

58. Peay, K.G., Kennedy, P.G. & Talbot, J.M. (2016). Dimensions of biodiversity in the Earth mycobiome. Nature Reviews Microbiology, 14, 434–447.

59. Peay, K.G., Russo, S.E., McGuire, K.L., Lim, Z., Chan, J.P., Tan, S., et al. (2015). Lack of host specificity leads to independent assortment of dipterocarps and ectomycorrhizal fungi across a soil fertility gradient. Ecology Letters, 18, 807–816.

60. Phillips, R.P., Brzostek, E. & Midgley, M.G. (2013). The mycorrhizal-associated nutrient economy: a new framework for predicting carbon-nutrient couplings in temperate forests. New Phytologist, 199, 41–51.

61. Pinheiro, J., Bates, D., DebRoy, S. & R Core Team. (2018). nlme: Linear and Nonlinear Mixed Effects Models. R package version 3.1–137.

62. Proctor, J., Anderson, J.M., Chai, P. & Vallack, H.W. (1983). Ecological Studies in Four Contrasting Lowland Rain Forests in Gunung Mulu National Park, Sarawak: I. Forest Environment, Structure and Floristics. The Journal of Ecology, 71, 237.

63. Read, D.J. & Perez-Moreno, J. (2003). Mycorrhizas and nutrient cycling in ecosystems - a journey towards relevance? New Phytologist, 157, 475–492.

64. Russo, S.E., Davies, S.J., King, D.A. & Tan, S. (2005). Soil-related performance variation and distributions of tree species in a Bornean rain forest. Journal of Ecology, 93, 879–889.

65. Terrer, C., Vicca, S., Hungate, B.A., Phillips, R.P. & Prentice, I.C. (2016). Mycorrhizal association as a primary control of the CO2 fertilization effect. Science, 353, 72–74.

66. Torti, S.D., Coley, P.D. & Janos, D.P. (1997). Vesicular-arbuscular mycorrhizae in two tropical monodominant trees. J. Trop. Ecol., 13, 623–629.

67. Torti, S.D., Coley, P.D. & Kursar, T.A. (2001). Causes and Consequences of Monodominance in Tropical Lowland Forests. Am Nat, 157, 141–153.

68. van der Heijden, M.G.A., Boller, T. & Wiemken, A. (1998). Different arbuscular mycorrhizal fungal species are potential determinants of plant community structure. Ecology.

69. van der Linde, S., Suz, L.M., Orme, C.D.L., Cox, F., Andreae, H., Asi, E., et al. (2018). Environment and host as large-scale controls of ectomycorrhizal fungi. Nature, 558, 243–248.

70. Zanne, A.E., Tank, D.C., Cornwell, W.K., Eastman, J.M., Smith, S.A., FitzJohn, R.G., et al. (2014). Three keys to the radiation of angiosperms into freezing environments. Nature, 1–10.

71. ZengPu, L. & JunRan, J. (1995). Antagonism between ectomycorrhizal fungi and plant pathogens. In: (eds. Brundrett, M.C., Dell, B., Malajczuk, N. & Mingqin, G.). Presented at the Mycorrhizas for Plantation Forestry in Asia, Kaiping, pp. 77–81.

